# A receptor for herbivore-associated molecular patterns mediates plant immunity

**DOI:** 10.1101/679803

**Authors:** Adam D. Steinbrenner, Maria Muñoz-Amatriaín, Jessica Montserrat Aguilar Venegas, Sassoum Lo, Da Shi, Nicholas Holton, Cyril Zipfel, Ruben Abagyan, Alisa Huffaker, Timothy J. Close, Eric A. Schmelz

## Abstract

Plant-herbivore interactions are ubiquitous across nature and drive major agricultural losses. Inducible defense responses triggered through immune recognition aid in host plant protection; however, specific ligand-receptor pairs mediating the initial perception of herbivory remain unknown. Plants in the subtribe Phaseolinae detect herbivore-associated peptides in caterpillar oral secretions and the defined ligands are proteolytic fragments of chloroplastic ATP synthase termed inceptins. Using forward genetic mapping of inceptin-induced responses, we identify a cowpea (*Vigna unguiculata*) leucine-rich repeat receptor-like protein as an inceptin receptor (INR) sufficient for elicitor-induced responses and enhanced defense against armyworms (*Spodoptera exigua*). INR defines a receptor by which plants perceive herbivore-associated molecular patterns (HAMPs) and expands the paradigm of surface immune recognition to attack with mandibles.

**One Sentence Summary:** A plant cell surface receptor directly perceives peptides associated with caterpillar herbivory.

## Main text

Herbivores defoliate and often devastate plants, making effective inducible defense responses critical for plant resilience (*1, 2*). To combat insect herbivores, plants respond to specific molecular patterns present in oral secretions (OS) and frass by amplifying wound-induced defenses and activating resistance (*3, 4*). Analogous immune responses to microbes and other pests are mediated by ligand-receptor interactions mediated by receptor kinases (RKs) or receptor-like proteins (RLPs) (*5*). While signaling and defensive outputs after herbivory are increasingly understood (*6*), receptors directly recognizing biochemically-defined herbivore-associated molecular patterns (HAMPs) have remained elusive (*7*).

Among defense-eliciting HAMPs, inceptins are a potent bioactive series of proteolytic fragments derived from the chloroplastic ATP synthase γ-subunit (cATPC) found in Lepidopteran larvae OS (*4, 8*). An abundant inceptin present during caterpillar herbivory on cowpea (*Vigna unguiculata*) is an 11 amino acid (AA) peptide, termed *Vu-*In (^+^ICDINGVCVDA^−^). The epitope is highly conserved among plant cATPC sequences; however, only species within the Phaseolinae subtribe respond to inceptins *(9).*

To identify inceptin receptor (INR) candidates, we examined plant response variation to both *Vu*-In and C-terminal truncated inceptin, termed *Vu*-In^−A^ (^+^ICDINGVCVD^−^), a less bioactive variant that uniquely accumulates in the OS of *Anticarsia gemmatalis,* a legume specialist herbivore (*10*). Anticipating an arms race pattern of evasion and re-establishment consistent with other elicitors (*11, 12*), we screened for cowpea germplasm with positive *Vu*-In^−A^-induced responses. Accessions Danila, Suvita, and Yacine displayed induced ethylene production after applying *Vu*-In^−A^ to wounded leaves, while other accessions failed to respond (Fig. 1A). Although responses to *Vu*-In^−A^ were quantitatively low compared to fully active *Vu*-In (Fig. 1A), we hypothesized that the existence of qualitative response variation to the weak elicitor variant could be mediated by INR genetic variation.

**Fig.1.**
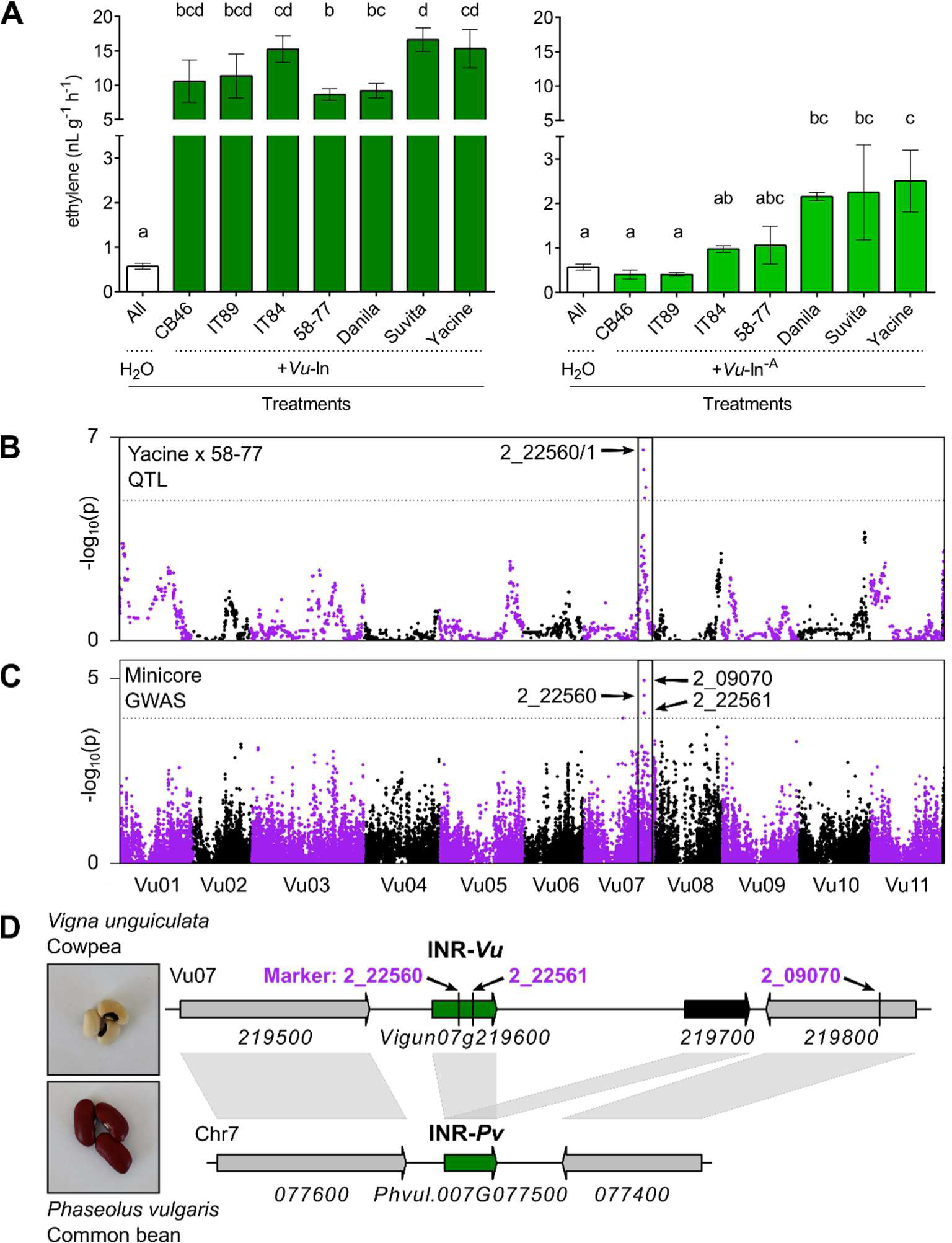
Cowpea sensitivity to *Vu*-In^-A^ associates with a single genetic locus. (A) Ethylene production of cowpea accessions after treatment with H_2_O or 1 μM inceptin peptides. Bars show means +/− SEM of n=3 replicate leaflets or all controls (n=21). Left, F=63.8, right, F=9.69, letters indicate significantly different means (Tukey HSD, α=0.05). (B) QTL statistics for ethylene production ratio, *Vu*-In^−A^/H_2_O, in the Yacine x 58-77 RIL population. Dotted line indicates significance cutoff using modified Bonferroni correction at α=0.05, –log_10_(p) > 4.84. (C) Manhattan plot of GWAS results for ethylene production ratio, *Vu*-In^−A^/H_2_O, in the panel of Minicore accessions. Dotted line indicates false discovery rate (FDR) significance cutoff at α =0.05, -log_10_(p) > 3.93. –log_10_ of the probability (p-values) is shown for 17638 (B) or 42686 (C) SNPs at respective physical coordinates in the cowpea genome (Lonardi et al. 2019). (D) Genomic region of Vu07 (positions 34,220,090-34,258,839) containing highly associated marker SNPs, and syntenic genes on common bean Chr7. Green and black filled arrows indicate RLP genes.

To map *INR,* we used a biparental population of recombinant inbred lines (RILs) derived from a cross between accessions Yacine (*Vu*-In^−A^ responsive) and 58-77 (*Vu*-In^−A^ unresponsive) (Fig. 1A) for QTL mapping, as well as a panel of 364 worldwide accessions belonging to the UC Riverside Minicore collection for a genome-wide association study (GWAS). *Vu*-In^−A^ elicited variable ethylene production across accessions (Fig. S1A-B, Table 1). Using QTL mapping and GWAS, we observed that *Vu*-In^−A^ responses strongly associated with a single genetic locus (Fig. 1B, 1C, Fig S2). The most highly-associated markers in both studies were SNPs 2_22560, 2_22561, and 2_09070 (*13*), spanning a 22-kb region on chromosome Vu07 (Fig. 1D, S1C). Both markers 2_22560 and 2_22561 fell within the receptor-encoding gene *Vigun07g219600* suggesting that this locus is associated with inceptin responsiveness in cowpea.

To examine function of the INR candidate, we transiently expressed *Vigun07g219600* from the reference accession IT97K-499-35 (*14*) in *Nicotiana benthamiana* and tested responsiveness to *Vu*-In. Hallmarks of receptor-mediated defense responses include a burst of reactive oxygen species (ROS) and ethylene production. As a positive control, transient expression of the *EF*-Tu receptor (EFR) (*15*) in *N. benthamiana* conferred responses to elf18 peptide (Fig. 2A). Expression of *Vigun07g219600* conferred *Vu-*In-inducible responses, in the form of ROS (Fig. 2B, 2C) and inducible ethylene production (Fig. 2D), indicating that *Vigun07g219600* encodes a functional INR.

**Fig. 2.**
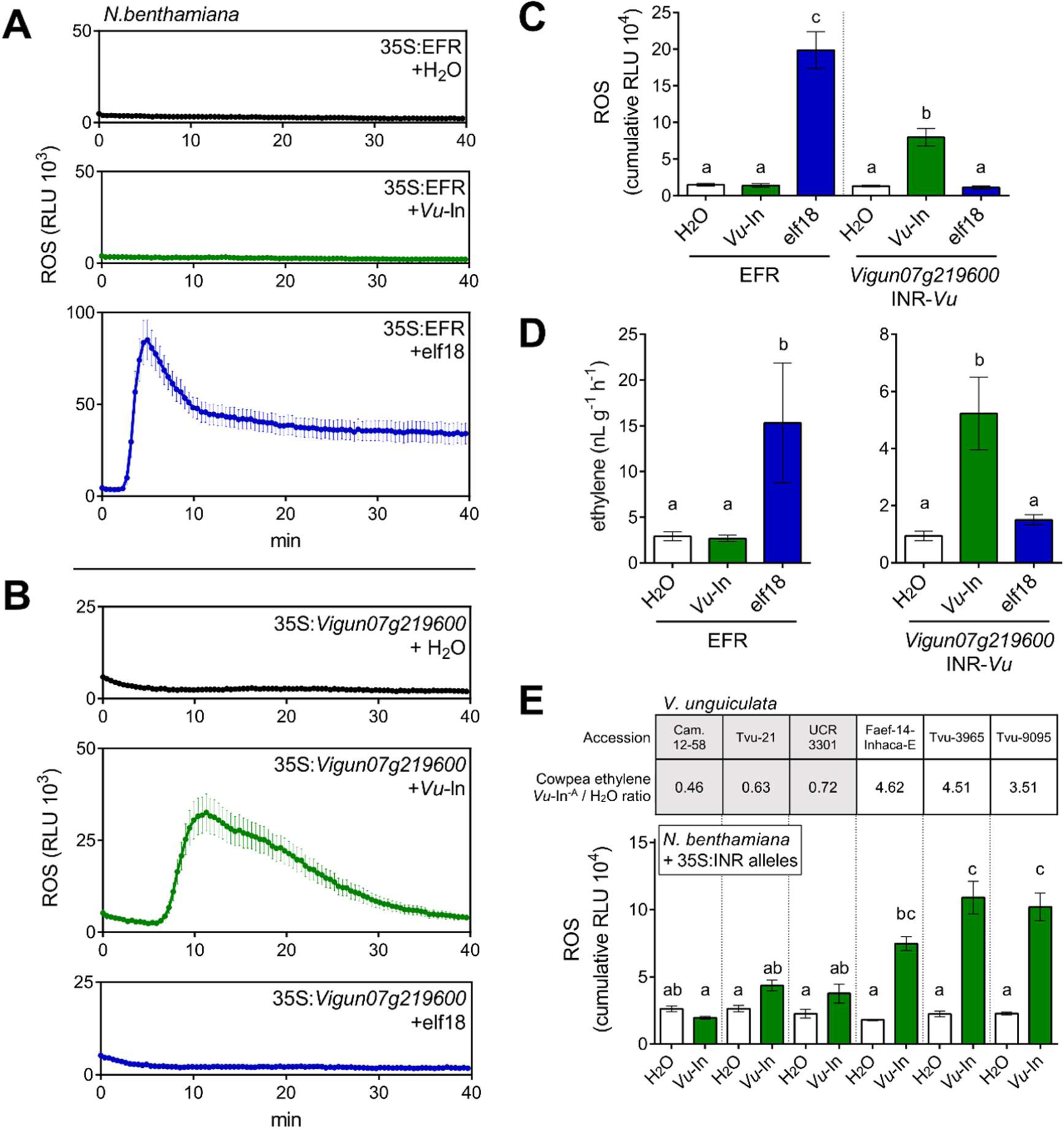
*Vigun07g219600*, Inceptin Receptor (INR), confers inceptin sensitivity in *N. benthamiana*. (A-C) Receptor-dependent reactive oxygen species (ROS) production in *N. benthamiana*. RLU, relative luminescence units, min, minutes after treatment with 1 μM peptide. Average is shown for n=8 leaf discs +/− SEM. (A) elf-18 receptor (EFR) expressed, (B) *Vigun07g219600* expressed. (C) Sum of RLU, 0-20 min. (D) Receptor-dependent ethylene production in *N. benthamiana* after treatment with water or 1 μM peptide. Bars show average of n=6 leaf discs +/− SEM. (E) Allelic variation in INR-dependent ROS in *N. benthamiana*. Bars show mean +/− SEM of n=8 leaf discs expressing alleles from indicated lines. Table indicates source accession for INR alleles. Letters indicate significantly different means (Tukey HSD, α=0.05).

To understand the basis of cowpea phenotypic variation for inceptin responsiveness, we cloned and expressed INR alleles from 6 cowpea accessions with differential *Vu*-In^−A^ responses. Only alleles from highly *Vu*-In^−A^ responsive accessions conferred significant *Vu*-In-induced ROS and ethylene production in *N. benthamiana*, thus mirroring *Vu*-In^−A^ response variation originally observed in cowpea (Fig. 2E, Fig S3). Interestingly, none of the tested alleles conferred *Vu*-In^−A^ responses in *N. benthamiana* (Fig. S3, S4). Given that all 364 cowpea accessions could respond to *Vu*-In (Table S1), natural variation in INR may thus specify an activation threshold for (1) the weak elicitor, *Vu*-In^−A^, in cowpea and (2) the robust elicitor, *Vu*-In, when expressed heterologously in *N. benthamiana.*

INR is a leucine-rich repeat (LRR)-RLP, a receptor class distinguished from LRR-RKs by lack of an intracellular kinase domain (*16*). INR contains 30 semi-regular LRRs preceding a transmembrane and short cytosolic domain (Fig. S5). The *INR* locus in cowpea contains a paralog *Vigun07g219700* (72% AA similarity) that is unable to confer *Vu*-In induced ethylene production when expressed in *N. benthamiana* (Fig. S6A). In contrast, two orthologs with >90% similarity, *Phvul.007G077500* (from *P. vulgaris*) and *Vradi08g18340* (from *V. radiata*), conferred *Vu-*In-induced ethylene production (Fig. S6B). In soybean (*Glycine max)*, the most closely related RLPs are four genes at the same syntenic locus, each with 73-76% similarity to cowpea INR-*Vu* (Fig. S6C). Neither of two tested soybean homologs (*Glyma.10G228000* or *Glyma.10G228100*) enabled *Vu-*In*-*induced ethylene production (Fig. S6B). Soybean plants are both unresponsive to inceptin and lack representation in a subclade of functional *INR* genes (Fig. S6C) (*9*). We conclude that *Phaseolus* and *Vigna*, both within the subtribe Phaseolinae, encode functional INRs.

RLPs from Arabidopsis and tomato have been demonstrated to bind peptide ligands (*17, 18*). To measure inceptin binding, we generated an N-terminal acridinium-tagged *Vu-*In conjugate. *N. benthamiana* expressing cowpea INR-*Vu* retained acridinium-*Vu-*In luminescent signal, while tissue expressing an unrelated receptor, EFR, did not (Fig. 3A). Furthermore, acridinium-*Vu*-In-derived luminescence was abolished by pre-incubating with excess unlabeled *Vu-*In (Fig. 3A). To better understand the receptor-ligand interaction, we constructed a homology model of INR based on the crystal structure of the LRR ectodomain of FLAGELLIN-SENSING 2 (FLS2) (*19*), and performed *Vu*-In docking simulations. The predicted lowest-energy conformations were ranked by peptide binding scores. In multiple low-energy conformations of *Vu*-In, the ligand acidic residue Asp10 showed a conserved binding position and conformation by interacting with both basic INR-*Vu* residues His495 and Arg497 (Fig. 3B). Consistent with critical binding activity, a previous Ala substitution study demonstrated that Asp10 was the only AA essential for *Vu*-In elicited ethylene production (*8*). To test the interaction, a receptor mutant was generated with Ala substitutions at His495/Arg497, which resulted in loss of both *Vu-*In binding activity and *Vu*-In-induced ROS production in *N. benthamiana* (Fig. 3A, 3B). His495 and Arg497 are highly conserved in *Phaseolus* and *Vigna* but not in soybean RLP homologs (Fig. S7). Our data supports INR-*Vu* binding to inceptin with key interactions specifically mediated by His495/Arg497.

**Fig. 3.**
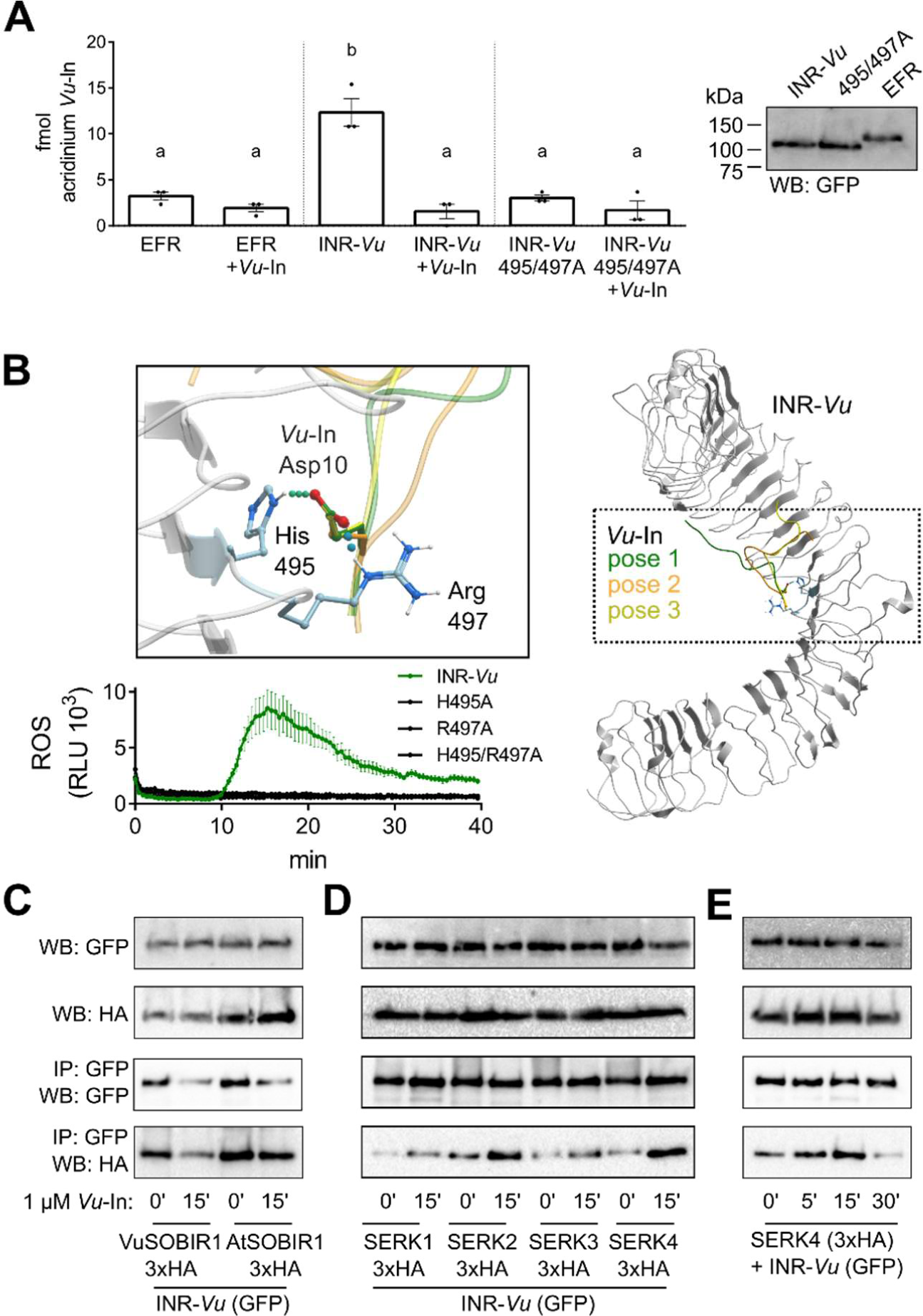
INR binds *Vu*-In peptide and associates with co-receptor RLKs. (A) INR-*Vu* but not EFR or mutant INR H495/R497A binds acridinium-*Vu*-In. Bars indicate mean +/− SEM (n=3 incubations) retained acridinium signal. Letters indicate significantly different means (Tukey HSD, α=0.05) (B) INR-*Vu* model in complex with predicted binding pose of the *Vu*-In peptide. Right, extracellular domain of INR-*Vu* was modeled by homology with the FLS2 ectodomain template (PDB code: 4mn8). The predicted top three *Vu*- In poses are shown. Left, close-up with *Vu*-In Asp10 residue sidechain shown interacting to INR-*Vu* His495 and Arg497 sidechains. Bottom, ROS burst (1 μM *Vu*-In) is shown for *N. benthamiana* leaf punches (n=8) expressing INR-*Vu* (green) or mutant variants (black). (C-F) INR-*Vu* co-immunoprecipitates (IP) Arabidopsis (At) and cowpea (Vu) homologs of SOBIR1 and SERK RLKs. C, VuSOBIR1 (*Vigun09g096400*) or AtSOBIR1 (*AT2G31880*) were co-expressed with INR-*Vu*. D-E, AtSERK1-4 were co-expressed with INR-*Vu.* All receptor (C-terminal GFP) and coreceptor (C-terminal 3xHA) combinations were expressed in *N. benthamiana.* Western blots (WB) were probed with either GFP or HA antibody. Representative results are shown from 2-3 independent experiments.

LRR-type surface receptors in plants typically associate with Somatic Embryogenesis Receptor Kinase (SERK) co-receptors for signal transduction (*20*). In addition, characterized RLPs constitutively associate with the adapter RK Suppressor of BIR1 (SOBIR1) (*21*). We tested if INR associates with Arabidopsis and cowpea orthologs of SOBIR1 by co-immunoprecipitation. Association of INR with both AtSOBIR1 and VuSOBIR1 (*Vigun09g096400*) was constitutive (Fig. 3C), while INR associated more strongly with AtSERK co-receptors after peptide treatment (Fig. 3D, 3E). Thus, INR associates with SERK and SOBIR1 receptor kinases in a manner similar to characterized LRR-RLPs.

To test if INR enhances anti-herbivore defenses in plants lacking native inceptin responses, we stably transformed *N. benthamiana* and *N. tabacum* with either 35S:INR-*Vu* or INR-*Pv* transgenes. Multiple independent transgenic lines expressing 35S:INR responded to *Vu*-In as measured by ethylene and a direct defense output, induced peroxidase activity (Fig. 4A,B, Fig. S9). We then challenged INR-expressing lines with second instar larvae of the generalist Lepidopteran herbivore, beet armyworm (*Spodoptera exigua*). Caterpillars consistently displayed 29-38% lower growth rates on transgenic lines than on wild-type (WT) when caged on single young seedlings of either *Nicotiana* species for 4 days (Fig. 4C, 4D). A similar magnitude of caterpillar growth reduction was previously observed after pretreatment of cowpea with *Vu-*In (*4*). Thus expression of INR-*Vu* or INR-*Pv* is sufficient to confer enhanced anti-herbivore defense responses.

**Fig. 4.**
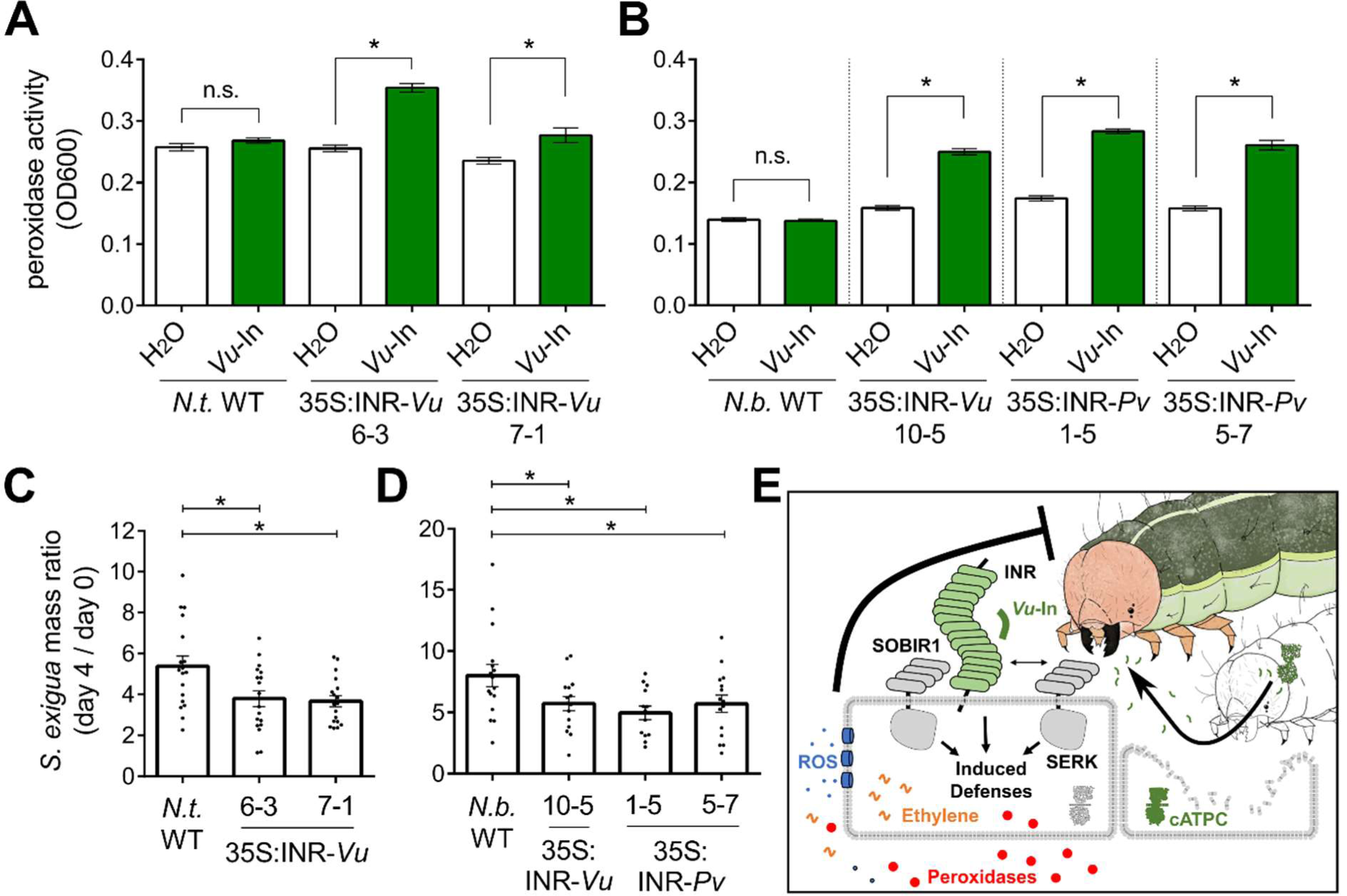
INR confers receptor activity and herbivore resistance to stable *Nicotiana* lines. (A-B) *Vu*-In induces peroxidase activity in stable INR transgenic lines of (A) *N. benthamiana (Nb)* and (B) *N. tabacum*. Bars show average +/− SEM of n=16-24 leaf discs after incubation with 1 μM inceptin. *, p<0.05 Student’s t-test. (C-D) Relative growth rates of beet armyworm (*Spodoptera* exigua) larvae reared on transgenic or WT lines of *N. benthamiana* (C) and *N. tabacum* (D). Bars show average +/− SEM of n=13-16 *N. benthamiana* plants or n=18 *N. tabacum* plants. For *N. benthamiana,* F=3.43, *N. tabacum*, F=5.93. * p<0.05, Fisher’s LSD. (E) Model for INR perception of *Vu-*In. INR associates with adapter and co-receptor kinases leading to burst of reactive oxygen species (ROS), ethylene, and peroxidases. Combined defense outputs negatively affect herbivore growth rate.

Dynamic plant defense responses to herbivory have been examined for nearly 50 years (*22*–*24*) and are induced by numerous insect associated elicitors or HAMPs (*3, 4*). In parallel with pathogen- and damage-associated molecular patterns (PAMPs and DAMPs), inceptin is detected by INR, an LRR-type surface receptor that establishes a HAMP-receptor pair. INR shares features of pattern recognition receptors, such as recent emergence in plant evolution and association with conserved signaling machinery (*25*). Despite similarities in mode of recognition, plant immunity to pathogens and herbivores is often characterized by downstream antagonistic interactions between defense hormones (*26*). INR and other diverse surface immune receptors will facilitate the comparison of signaling tradeoffs against a wide breadth of pest classes. A defined ligand-receptor pair for plant-herbivore interactions represents a step forward in both understanding and controlling plant immunity to diverse biotic pressures.

## Acknowledgments

We thank Harley Riggleman, Nile J. Hodges, and Marlo Hall for technical and artistic assistance. We thank Ted Turlings, Yezhang Ding, Elly Poretsky, Keini Dressano, and Philipp Weckwerth for advice and discussions.

## Data and Materials Availability

INR allele sequences are available at Genbank (submission #2217536)

