## Supplementary Materials for "A receptor for herbivore-associated molecular patterns mediates plant immunity"

1. Supplementary Methods
2. Supplementary Figures S1-S8
3. Supplementary Tables 1-2
4. References 27-43

### 1. Supplementary Methods

#### Cowpea materials and SNP genotyping

Two populations were used for mapping of Vu-In$^{-A}$ response: a bi-parental recombinant inbred line (RIL) population of 109 lines derived from a cross between Yacine and 58-77 that was developed previously (*27*), and a set of 364 cowpea accessions representing worldwide diversity of cultivated cowpea (M. Muñoz-Amatriaín, S. Lo, and T. J. Close, personal communication). Both populations were genotyped with the Cowpea iSelect Consortium Array containing 51,128 SNPs (*12*) at the University of Southern California Molecular Genomics Core facility (Los Angeles, CA, USA). SNPs were called using GenomeStudio software (Illumina, Inc., San Diego, CA, USA) with the custom file from Muñoz-Amatriaín et al. (*12*). Data curation was performed by removing SNPs with more than 20% missing or heterozygous calls.

#### Linkage and QTL mapping

RILs with high heterozygosity and those carrying non-parental alleles were eliminated prior to mapping. This resulted in the removal of nine RILs. Only 17,638 SNPs that were polymorphic in both the parents and the RIL population, and that had minor allele frequencies (MAFs) >0.20 were used for linkage mapping. MSTmap (*28*) (http://www.mstmap.org/) was used for genetic map construction, with the following parameters: grouping LOD criteria = 10; population type = DH (doubled haploid); no mapping size threshold = 2; no mapping distance threshold: 10 cM; try to detect genotyping errors = no; and genetic mapping function = kosambi. The linkage groups were numbered and oriented according to cowpea pseudomolecules (*29*). Since the DH function inflated the cM distance for a RIL population, cM distances were divided by two to correct for the extra round of effective recombination occurring in a RIL population compared to a DH population.

QTL analysis was performed using a linear mixed model described by Xu (*30*) and implemented in R by Lo et al.(*31*). A modified Bonferroni correction ($\alpha$ =0.05) that uses the "effective number of markers or effective degrees of freedom" instead the total number of SNPs as a denominator was used to set the genome-wide critical value, as in Lo et al. (*31*). This set the significance cut-off to a $-\log_{10}$(p-value) of 4.84 for mapping *Vu*-In$^{-A}$ response.

**Genome-wide association studies (GWAS)**

GWAS was performed in the panel of UCR Minicore accessions using the mixed linear model (MLM) function (*32*) implemented in TASSEL v5 (www.maizegenetics.net/tassel), with a principal component analysis (5 principal components) accounting for population structure in the dataset and a kinship matrix correcting for genetic relatedness between accessions. A total of 42,686 SNPs with MAF>0.05 were used for GWAS. SNPs were ordered based on their physical position in the cowpea reference genome (*29*). A false discovery rate ($\alpha$ =0.05) was used for multiple testing correction of the GWAS results, which set the significance threshold to a $-\log_{10}$(p-value) of 3.93 for mapping *Vu*-In$^{-A}$ response.

**Peptide-induced ethylene production**

Inceptin peptides based on *Vigna unguiculata* cATPC synthase sequence, *Vu*-In (ICDINGVCVDA) and *Vu*-In$^{-A}$ (ICDINGVCVD), were ordered from Genscript and reconstituted in water.

Induced ethylene accumulation in cowpea was measured in first fully-extended trifoliate leaves of 3-week old greenhouse-grown seedlings. Leaflets were lightly scratch-wounded in each corner with a fresh razor blade to remove cuticle over an area of 1 $cm^2$, and 10 µL of water with or without peptide was equally spread over the 4 wounds with pipette tip. After 1 hour, leaflets were excised and placed in sealed tubes for 1 hour before headspace sampling. Ethylene was measured by GC-FID (Supelco 13018-U, 80/100 Hayesep Q) and quantified using a standard curve.

For QTL mapping, experimental design was as follows: 2 plants of each of 85 successfully germinated recombinant inbred lines (RILs) were treated with water or Vu-In$^{-A}$ as described above, on paired leaflets of either trifoliate or primary leaves. The ratio of ethylene production per gram of tissue (Vu-In$^{-A}$ : $H_2O$) was calculated for each individual pair of leaflets, and the log-corrected average of the 4 pairs was used for QTL mapping. For GWAS, experimental design was similar except 1 trifoliate leaf of a single seedling of each of 364 lines were treated.

For ethylene assays in *N.benthamiana*, most recently fully extended leaf of 4-week old plants was infiltrated with blunt syringe, patted dry with paper towel, and 4 leaf discs within infiltrated area were immediately excised with cork borer and sealed in tubes. Headspace ethylene was measured after 3 hours of accumulation.

**Molecular Cloning, Transient and Stable Expression**

Full length cDNA sequences of all described receptors and co-receptors were PCR amplified using primers listed in Table S2 from 5' SMARTer RACE cDNA libraries (Takara Biosciences). For the highly similar soybean genes *Glyma.10G228000* and *Glyma.10G228100*, a larger fragment was subcloned using primers with local homology for flanking sequences on chromosome 10, then gene-specific primers were used to amplify coding sequence. Amplicons were inserted using Gateway technology (Invitrogen) into plant expression vectors pEarleyGate103 (*33*) for C-terminal GFP or pGWB414 (*34*) for C-terminal 3xHA tag. Constructs were electroporated into *Agrobacterium tumefaciens* strain GV3101 (pMP90) (*35*). Agrobacterium strains for expression of individual constructs were induced with 150 µM acetosyringone in 10 mM MES pH 5.6, 10 mM $MgCl_2$ and infiltrated into *N.benthamiana* leaves at OD600 of 0.45.

Transgenic lines of *Nicotiana benthamiana* and *Nicotiana tabacum* var. SR1 were obtained from the UC Davis Plant Transformation Facility using GV3101 strains (pEG103) with INR-*Vu* or INR-*Pv* (C-terminal GFP) inserts. T1 lines with segregation of herbicide resistance consistent with single transgene insertions were selfed and homozygous T2 lines selected.

**Homology modeling of INR-*Vu* and Docking of *Vu*-In**

The crystal structure of LRR ectodomain of FLS2 (PDB ID: 4MN8) was used as the template. Sequence of INR-*Vu* was aligned with sequence of the template through the zero end-gap global alignment (ZEGA) method with the Gonnet comparison matrix (*36*, *37*). The penalty of gap opening and extension were set as 2.4 and 0.15, respectively. Based on the alignment and template structure, a homology model of INR-Vu was built with the homology modeling tool and default parameters in ICM-Pro (*38*). All side chains and insertions/deletions were refined via a biased probability Monte Carlo (BPMC) sampling (*39*)

The co-crystallized 22-amino-acid peptide in the crystal structure template (PDB ID: 4MN8) was used to define the docking region. A set of potential maps were generated for the docking region on a 0.5 Å 3D grid, containing: (i) van der Waals interaction; (ii) electrostatic interaction; (iii) hydrogen bond; and (iv) hydrophobic potential grids. With potential maps, docking and scoring of *Vu*-In was performed using a stochastic global energy optimization procedure in internal coordinates implemented in the ICM-Pro v3.8-6a (*40*), described as the following steps. 1) The *Vu*-In peptide was sampled with the implicit solvation model to generate a series of starting conformations via BPMC method, and each starting conformation was placed into the docking region with four principal orientations. 2) The *Vu*-In was sampled in the pre-calculated potential maps through BPMC sampling to optimize its positional and internal variables. 3) After sampling, 10 top ranking conformations were re-scored with ICM full atom scoring function and conformations were re-sorted by the docking score.

**Acridinium-labeled peptide**

Acridinium-labeled *Vu*-In (acri-In) was synthesized using N-hydroxy-succinimidyl acridinium ester (Cayman Chemical, Ann Arbor USA), quenched with Tris pH 8, and labeled peptide was separated by HPLC (Agilent Poroshell 120 EC-C18) by tracking absorbance at 372 nM. Final labeled peptide concentration was determined using a standard curve of 372 nm absorbance of NHS-acridinium standard.

Binding assays were performed according to (*41*) with modification. Pre-ground *N.benthamiana* tissue (48 hpi Agrobacterium), was vortexed in binding buffer (50 mM Tris pH 7.5, 150 mM NaCl, 1 mM PMSF, 1x Roche Protease Inhibitor Cocktail), spun 5' at 20,000 rcf, and pellet was resuspended in binding buffer at 2 mL/g. Tissue pellet resuspension was divided by aliquoting 80 µL per replicate tube. Four microliters of either water or excess competitor peptide was added, to a final concentration of 40 µM, and tissue was pre-incubated on ice for 2 hr. Acridinium-labeled peptide in binding buffer was added to a final concentration of 200 nM with or without additional competitor peptide (final concentration 80 µM in 100 µL volume). After 20 min incubation on ice with occasional mixing, pellet was washed three times with binding buffer and resuspended in 100 µL 5 mM citric acid. Pellet-retained luminescence was measured using a Biotek Synergy H2 Multimode plate reader by injecting 100 µL trigger buffer (0.2 N NaOH, 0.1% $H_2O_2$) and reading 10s luminescence, and a standard curve was used to determine retained peptide concentration.

**Co-immunoprecipitation**

Forty-eight hours post-infiltration with Agrobacteria, *N.benthamiana* leaves expressing both INR or EFR (C-terminal GFP) and SOBIR1 or SERKs (C-terminal 3xHA) were infiltrated with peptide solutions and harvested on liquid $N_2$ at specified timepoints. Tissue was ground on $N_2$-chilled mortar and pestle and homogenized in 2 mL/g extraction buffer (50 mM Tris pH 7.5, 150 mM NaCl, 1% Nonidet P-40, 10 mM DTT, 1x Roche Protease Inhibitor) then cleared by centrifugation (30m, 21 krcf). Supernatant was incubated with 10 µL GFP-Trap A beads (Chromotek) by end-over-end mixing at 4°C and eluted in 30 uL of Laemli buffer (95°C, 5 min).

**Reactive Oxygen Species (ROS) Production and Peroxidase Activity Assays**

Twenty-four hours post-infiltrated with Agrobacteria, leaf punches were taken with a biopsy punch and floated in water. After overnight incubation, ROS production was measured using luminol-horseradish peroxidase (HRP) over 40m as described in (*42*) using a Biotek Synergy H2 Multimode plate reader.

Peptide-induced peroxidase activity was measured as described by Mott *et al. (43)* with the following modifications. Leaf discs were taken from fully extended leaves of 4-week old *Nicotiana* seedlings, washed for 1 hour in ½ Murashige Skoog salt (4.4 g/L), and incubated overnight in ½ MS with 1 µM peptide prior to 5-aminosalicylic acid reaction with resulting solution as described.

**Herbivory Assays**

Beet armyworm (*Spodoptera exigua*) larvae were ordered from Benzon (Carlisle, PA) as first instars, incubated at 28°C for 48 h, then newly molted second instars were pre-weighed and placed on individual 3-week old plants. Each plant was contained with individual larvae using PVC cages tamped around surrounding soil material. *Nicotiana* with caterpillars were kept in a 16/8 light/dark cycle at 28°C. After 4 days, larvae were weighed and relative growth rate was calculated.

### 3. Supplementary Figures S1-S8

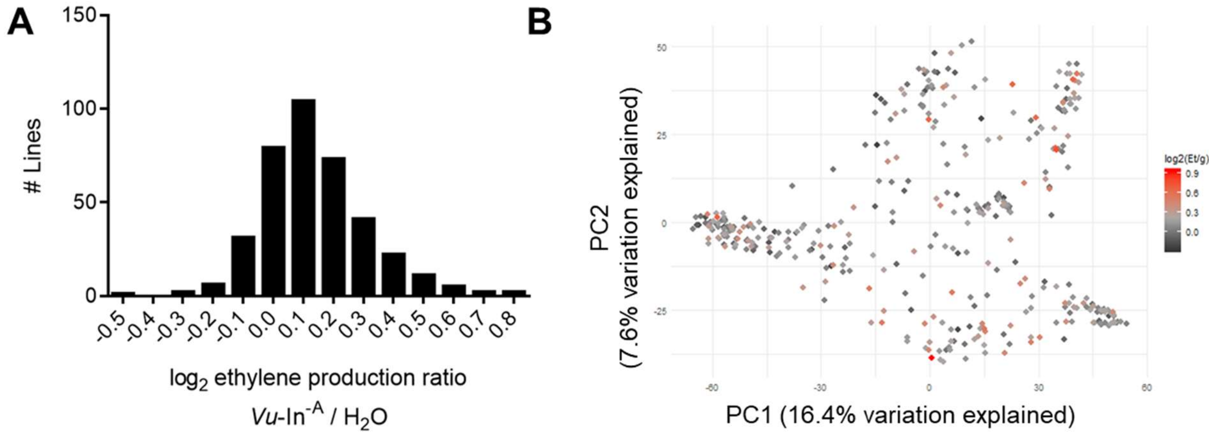


C

QTL

| Marker | Chr | position | $-\log_{10}(p)$ |
|---|---|---|---|
| 2_22560 | 7 | 34235351 | 6.592 |
| 2_22561 | 7 | 34236060 | 6.592 |
| 2_09594 | 7 | 34212130 | 6.592 |
| 2_09596 | 7 | 34216714 | 6.592 |
| 2_51758 | 7 | 34630814 | 5.921 |
| 2_06291 | 7 | 35676057 | 5.306 |
| 2_10029 | 7 | 35683365 | 5.306 |
| 2_08752 | 7 | 35683365 | 5.306 |
| 2_16606 | 7 | 35086036 | 4.943 |
| 2_49081 | 7 | 35086036 | 4.943 |

GWAS

| Marker | Chr | position | $-\log_{10}(p)$ |
|---|---|---|---|
| 2_09070 | 7 | 34256864 | 4.959 |
| 2_22560 | 7 | 34235351 | 4.548 |
| 2_22561 | 7 | 34236060 | 4.076 |
| 2_02375 | 7 | 22014760 | 3.932 |

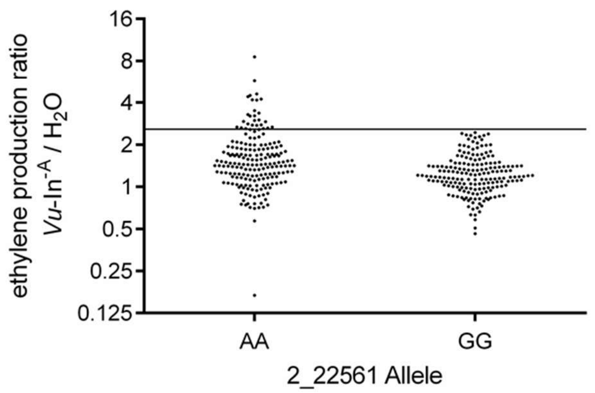


**Fig. S1, Supplement for Fig 1, Cowpea elicitation by Vu-In-A associates with a single genetic locus.** (A) Cowpea accessions vary in *Vu*-In$^{-A}$ induced ethylene. Lines were binned into centered groups based on $\log_2$-corrected ratio of ethylene production, *Vu*-In$^{-A}$ / $H_2O$. (B) Genetically diverse cowpea accessions respond to *Vu*-In$^{-A}$. UCR Minicore lines separated by principal component (PC) distance and colored by $\log_2$-corrected ratio of ethylene production, *Vu*-In$^{-A}$ / $H_2O$ (C) List of all significantly associated SNPs (FDR < 0.05) and physical positions from QTL mapping ($-\log_{10}(p) > 4.84$) and GWAS ($-\log(p) > 3.93$). See Fig. 1B and 1C for summary statistics for all markers. (D) Cowpea lines with 2_22561 allele AA respond more strongly to *Vu*-In$^{-A}$. Points indicate ratio of induced ethylene, *Vu*-In$^{-A}$ to water, for 364 individual accessions from the UCR Minicore with either AA or GG at marker SNP 2_22561.

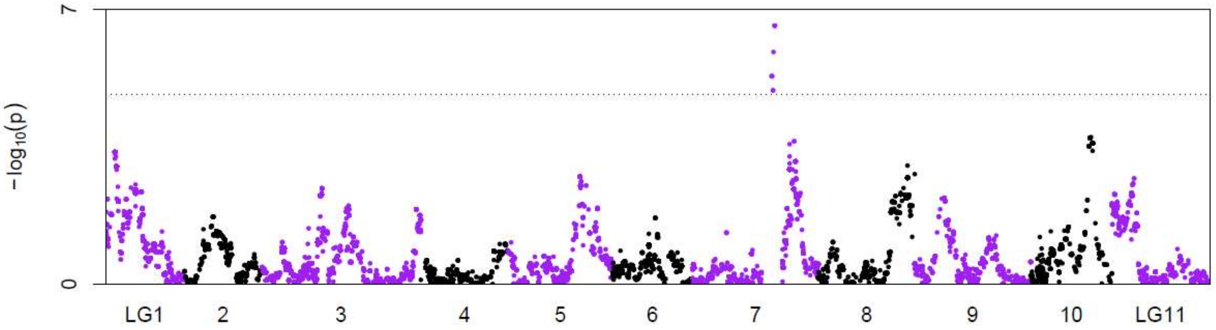


**Fig. S2, Statistics of Vu-In$^{-A}$ response QTL mapping on the cowpea genetic map from the Yacine x 58-77 RIL population.** Statistics as in Fig. 1B, but with positions indicating cM in 11 linkage groups (LG).

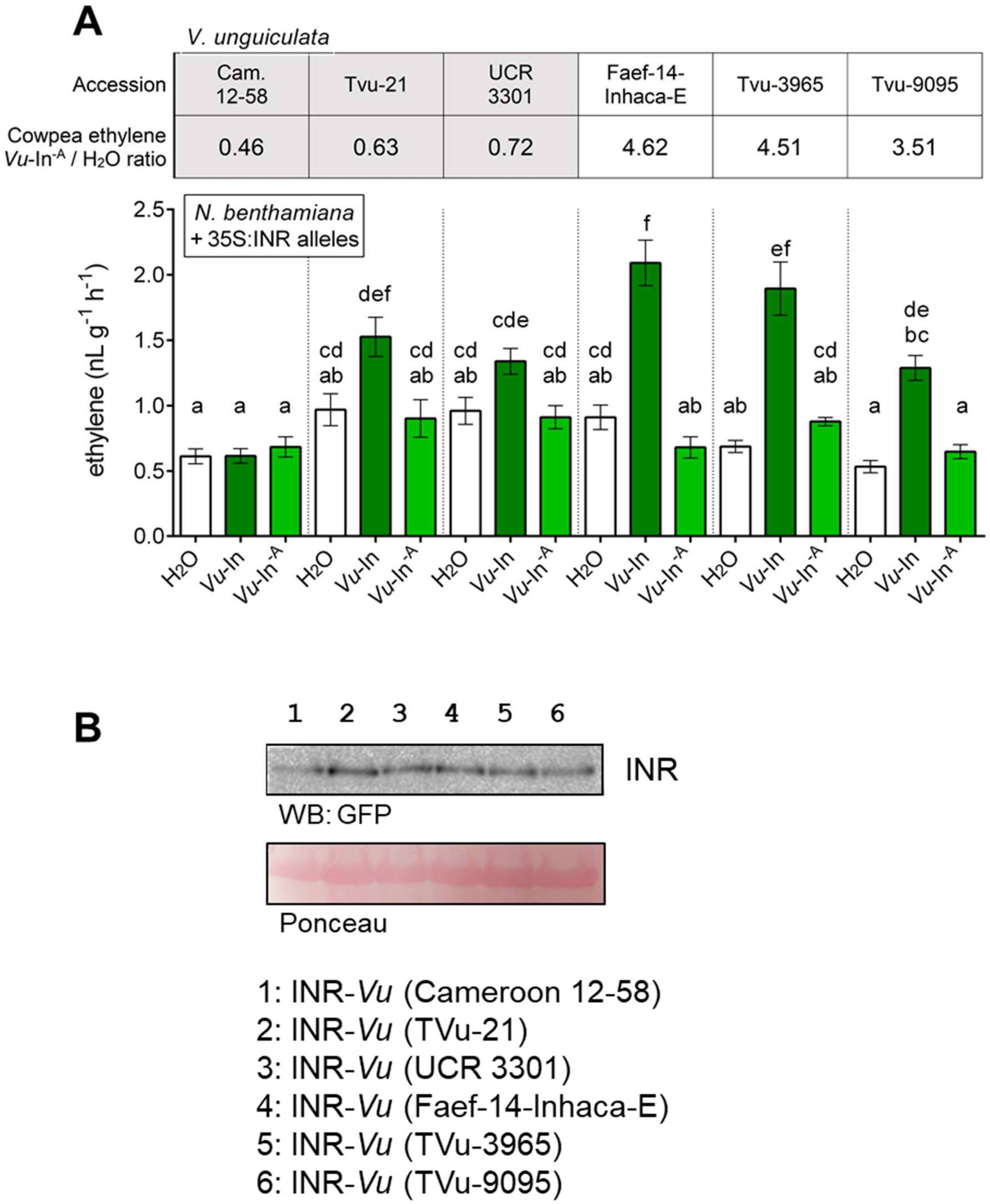


**Fig. S3, Cowpea INR allelic variation corresponds with function in *N.benthamiana.*** (A) *Vu*-In induced ROS production varies by allele. INR-*Vu* alleles from six cowpea varieties were expressed in *N.benthamiana* leaves, leaf punches taken at 24 hpi were treated with water or peptide, and total ROS produced over 25 min was summed. Bars show average sum of n=6 leaf discs +/- SEM, Letters indicate significantly different means (Tukey HSD, $\alpha$=0.05). (B) Protein expression of different INR-*Vu* alleles expressed in *N.benthamiana* and harvested 48 hours post-inoculation (hpi).

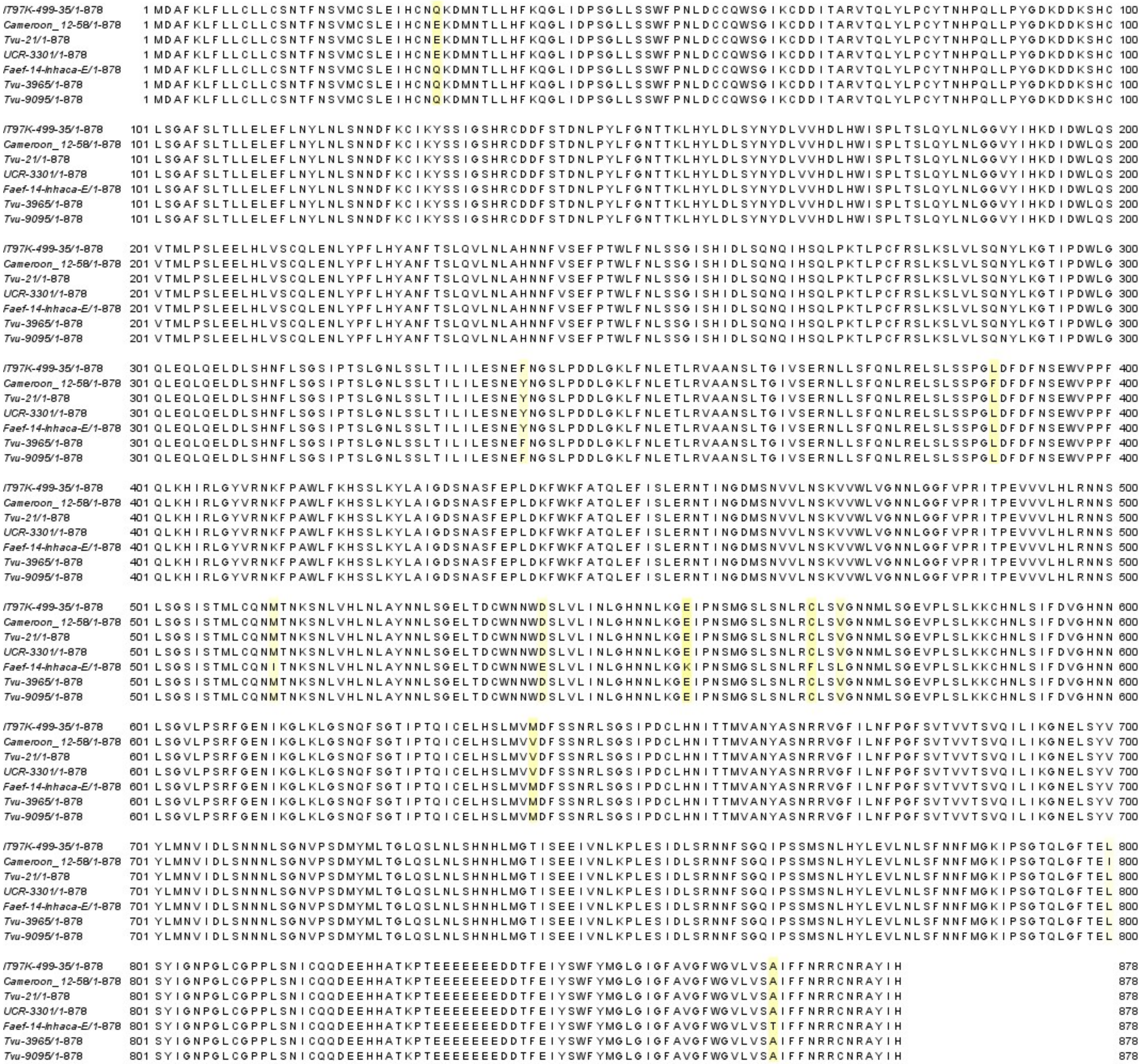

**Fig. S4. Amino acid alignment of INR alleles from reference cowpea (IT97K-499-35) or cowpea lines from UCR Minicore.** Polymorphic amino acids are highlighted in yellow. Alignment was visualized using Jalview.

```
Signal peptide        MDAFKLFLLCLLCSNTFNSVMC

N-terminal cap        SLEIHCNQKDMNTLLHFKQGLIDPSGLLSS
                      WFPNLDCCQWSGIKCDDITA

1  LRRs                         RVTQLYLPCYTNHP    QLLP
2                               YGDKDDKSHCLSGAF
3                             SLTLLELEFLNYLNLSNNDFKCIK
4                                   YSSIGSHRCDDFS TDNLP
5                     YLFGNTTKLHYLDLSYNYDLVVHDL HWISP
6                                LTSLQYLNLGGVYI HKDID
7                         WLQSVTMLPSLEELHLVSCQL ENLYP
8                          FLHYANFTSLQVLNLAHNNF VSEFP
9                          TWLFNLSSGISHIDLSQNQI HSQLP
10                          KTLPCFRSLKSLVLSQNYL KGTIP
11                          DWLGQLEQLQELDLSHNFL SGSIP
12                          TSLGNLSSLTILILESNEF NGSLP
13                          DDLGKLFNLETLRVAANSL
14                         TGIVSERNLLSFQNLRELSL   SSP
15                      GLDFDFNSEWVPPFQLKHIRLGYVRNKFP
16                          AWLFKHSSLKYLAIGDSNAS  FEP
17                        LDKFWKFATQLEFISLERNTI NG
18                          DMSNVVLNSKVVWLVGNNL GGFVP
19                              RITPEVVVLHLRNNSLSGSIS
20                      TMLCQNMTNKSNLVHLNLAYNNL SGELT
21                          DCWNNWDSLVLINLGHNNL KGEIP
22                          NSMGSLSNlLRCLSVGNNMLSGEVP
23                          LSLKKCHNLSIFDVGHNNL SGVLP
24                            SRFGENIKGLKLGSNQF SGTIP
25                          TQICELHSLMVMDFSSNRL SGSIP
26                        DCLHNITTMVANYASNRRVGF ILNFP
27                              GFSVTVVTSVQILI
28                        KGNELSYVYLMNVIDLSNNNL SGNVP
29                          SDMYMLTGLQSLNLSHNHL MGTIS
30                          EEIVNLKPLESIDLSRNNF SGQIP
31                          SSMSNLHYLEVLNLSFNNF MGKIP

C-terminal capping    SGTQLGFTELSYIGNPGLCGPPLSNICQQ

Acidic                DEEHHATKPTEEEEEEEDDTFEIYS

Transmembrane         WFYMGLGIGFAVGFWGVLVSAIFF

Cytosolic             NRRCNRAYIH
```

**Fig. S5, Amino acid sequence of INR-*Vu* and subdomains.** Conserved leucine, asparagine, and proline residues in the LRR motif are highlighted. Putative peptide binding residues are highlighted in green. Polymorphic marker SNP 2_22561 (allele either AA=Met or GG=Val) is highlighted in red.

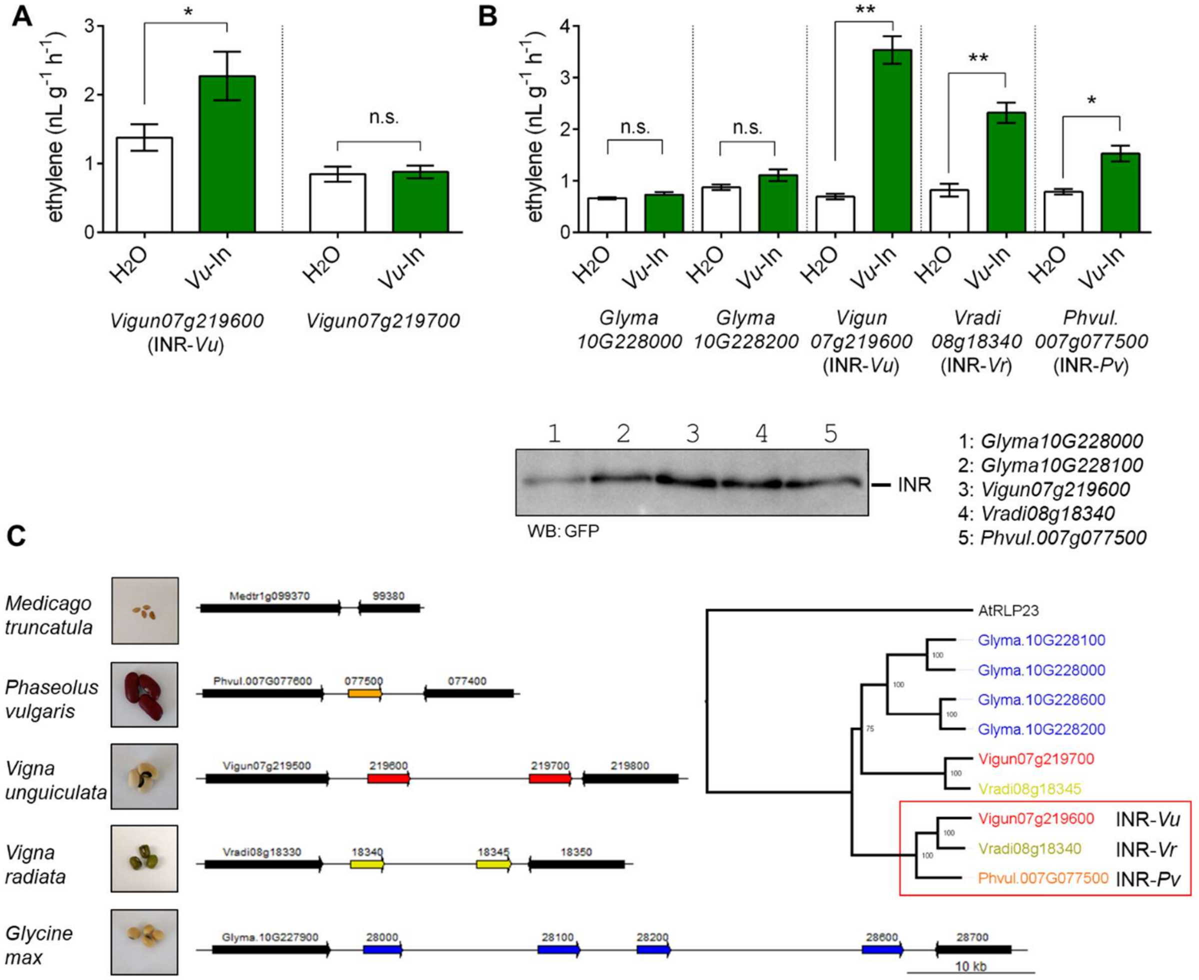


**Fig. S6, Orthologous RLPs within the Phaseolinae form an INR clade.** (A) *V. unguiculata* paralogous receptor *Vigun07g219700* is insufficient to confer *Vu*-In sensitivity. Genes were transiently expressed in *N. benthamiana* and leaves were infiltrated with water or 1 µM *Vu*-In and ethylene accumulation was measured. Bars indicate average of n=6 leaves with SEM, * paired t-test p<0.05. (B) INR activity of homologous leucine-rich repeat receptor-like proteins (LRR-RLPs). Receptors were expressed with C-terminal GFP tag in *N. benthamiana* and treated at 48 hpi with 1 uM *Vu-In* peptide for ethylene assay. Bars indicate average of n=6 leaves with SEM, * paired t-test p<0.05. Western blot analysis of RLP receptor stable expression confirmed accumulation of all receptor protein products. (C) Syntenic *INR* locus in different legume species, from top to bottom *Medicago truncatula, P. vulgaris, V. unguiculata, V. radiata,* and *G. max*. Colored arrows indicate LRR-RLP. Right, BioNJ tree was generated in Seaview based on outcome of 500 bootstraped replicates, with AtRLP23 (AT2G32680) as outgroup. Red box indicates INR clade containing all homologs sufficient to confer inceptin responses in *N. benthamiana* assays.

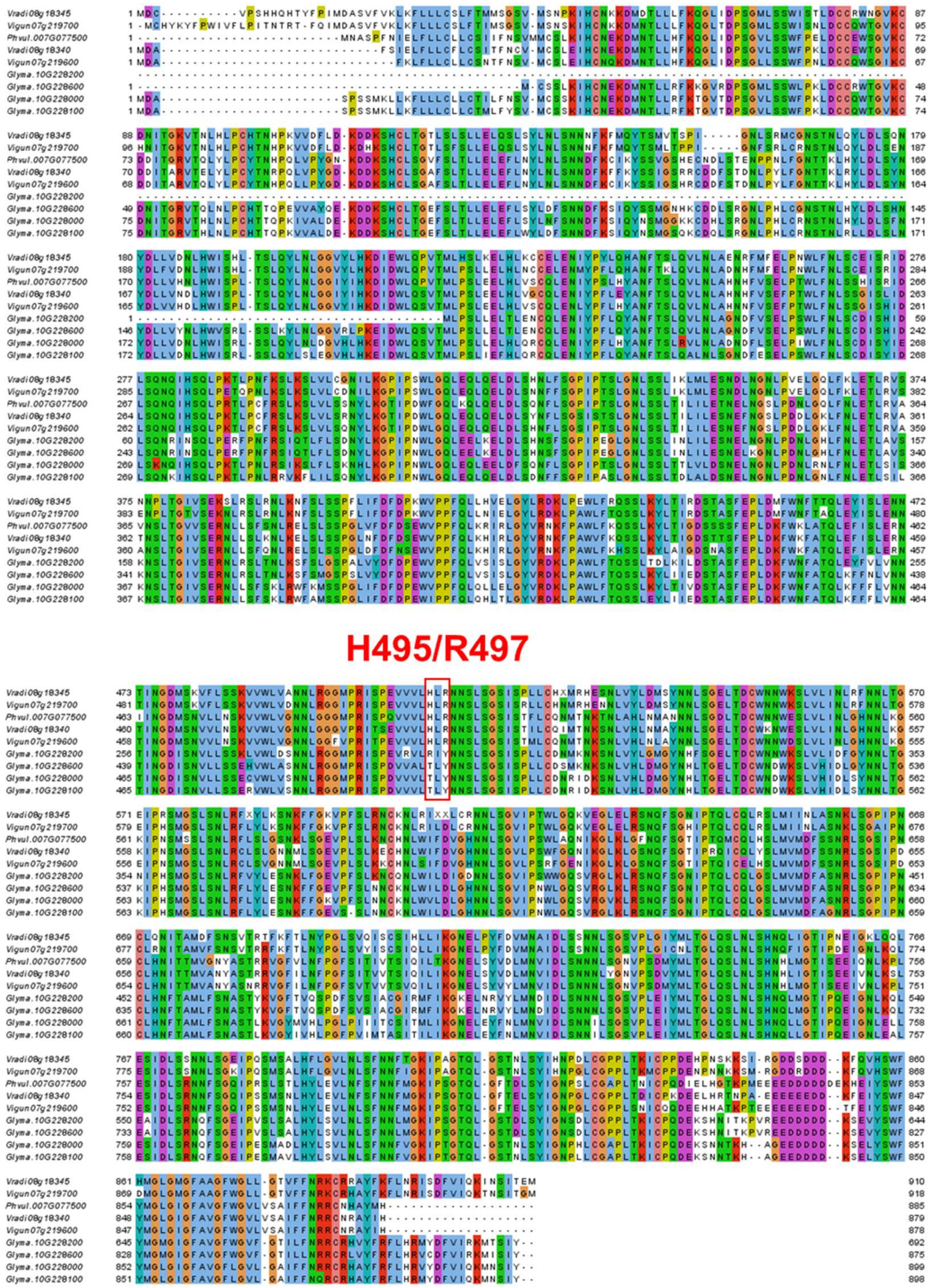


**Fig. S7, Multiple sequence alignment of INR homologs.** Coding sequences were retrieved, aligned with MUSCLE, and visualized in Jalview. Putative peptide binding site is highlighted with red rectangle

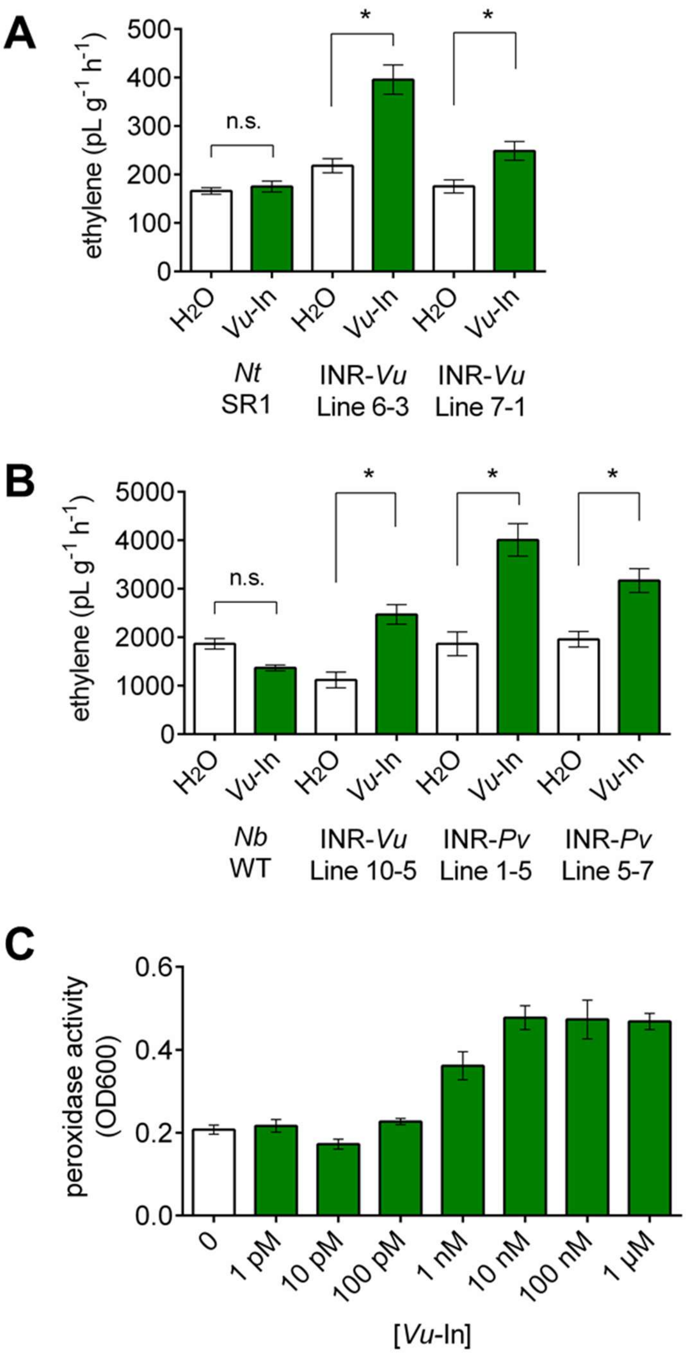


**Fig. S8, *Vu*-In responses in transgenic *Nicotiana*.** (A) Inducible ethylene (Et) biosynthesis in INR transgenic *Nicotiana tabacum* T2 lines. Individual leaves were infiltrated with H2O or 1 µM *Vu*-In peptide. Bars show average +/- SEM, n=10 leaves. (B) Inducible ethylene (Et) biosynthesis in INR transgenic *Nicotiana benthamiana* T2 lines. Individual leaves were infiltrated with $H_2O$ or 1 µM *Vu*-In peptide. Bars show average +/- SEM, n=3 leaves. *, $p<0.05$ Student's paired t-test, n.s. not significant.

### 2. Supplementary Tables 1-2

| Accession | *Vu*-In$^{-A}$ : $H_2O$ | *Vu*-In : $H_2O$ | Accession | *Vu*-In$^{-A}$ : $H_2O$ | *Vu*-In : $H_2O$ | Accession | *Vu*-In$^{-A}$ : $H_2O$ | *Vu*-In : $H_2O$ | Accession | *Vu*-In$^{-A}$ : $H_2O$ | *Vu*-In : $H_2O$ | Accession | *Vu*-In$^{-A}$ : $H_2O$ | *Vu*-In : $H_2O$ | Accession | *Vu*-In$^{-A}$ : $H_2O$ | *Vu*-In : $H_2O$ |
|---|---|---|---|---|---|---|---|---|---|---|---|---|---|---|---|---|---|
| 1393-1-2-3(-) | 1.05 | 10.74 | IT98D-1399 | 1.47 | 10.67 | TVu-13463 | 0.81 | 38.40 | TVu-1637 | 1.99 | 22.00 | TVu-6356 | 1.70 | 44.94 | TVu-9508 | 1.16 | 18.37 |
| 24-125-B-1 | 1.26 | 22.96 | IT98K-1111-1 | 1.13 | 11.38 | TVu-13485 | 1.30 | 44.27 | TVu-16403 | 1.02 | 43.08 | TVu-6365 | 2.11 | 24.31 | TVu-9516 | 1.94 | 43.00 |
| 524B | 1.33 | 8.57 | IT98K-128-4 | 1.13 | 26.35 | TVu-13778 | 1.69 | 78.22 | TVu-16408 | 1.09 | 7.23 | TVu-6439 | 1.40 | 89.60 | TVu-9522 | 1.66 | 42.80 |
| 58-53 | 0.94 | 26.35 | IT98K-205-8 | 1.40 | 6.36 | TVu-13939 | 1.24 | 17.86 | TVu-16449 | 0.81 | 37.00 | TVu-6464 | 2.00 | 14.55 | TVu-9556 | 1.18 | 22.77 |
| 58-57 | 1.53 | 88.67 | IT98K-428-2 | 1.25 | 38.86 | TVu-13950 | 1.28 | 17.50 | TVu-16465 | 1.69 | 5.07 | TVu-6477 | 1.25 | 22.22 | TVu-9557 | 1.16 | 54.80 |
| Apagbaala | 1.52 | 27.83 | IT98K-498-1 | 0.80 | 27.43 | TVu-13958 | 1.35 | 7.50 | TVu-16504 | 0.87 | 7.22 | TVu-6493 | 2.77 | 22.15 | TVu-9620 | 1.85 | 24.53 |
| Bambey_21 | 1.38 | 17.00 | IT98K-555-1 | 2.18 | 10.71 | TVu-13965 | 1.98 | 25.85 | TVu-16505 | 1.99 | 21.05 | TVu-6641 | 1.07 | 55.11 | TVu-9651 | 1.05 | 32.00 |
| Bun_22_Ghana | 1.41 | 12.00 | IT99K-124-5 | 1.32 | 11.36 | TVu-13968 | 1.11 | 40.00 | TVu-16521 | 1.02 | 45.54 | TVu-6642 | 1.45 | 23.35 | TVu-969 | 1.50 | 32.48 |
| CB27 | 1.28 | 10.00 | Ife_Brown | 1.03 | 34.37 | TVu-13979 | 1.24 | 27.90 | TVu-1715 | 0.70 | 26.84 | TVu-6643 | 1.55 | 42.07 | TVu-972 | 1.73 | 21.57 |
| CB3 | 1.07 | 26.50 | Iron_Clay_R | 1.21 | 62.55 | TVu-14172 | 1.20 | 24.22 | TVu-1727 | 0.86 | 17.45 | TVu-6644 | 1.68 | 16.98 | TVu-9749 | 1.67 | 37.50 |
| CB46 | 1.28 | 62.93 | KVx-396-4-5-20 | 0.17 | 11.88 | TVu-14190 | 1.70 | 27.00 | TVu-1775 | 1.40 | 43.89 | TVu-6663 | 0.86 | 59.20 | TVu-9801 | 1.28 | 49.23 |
| CB50 | 1.13 | 26.67 | KVx-403-P-20-T | 0.90 | 50.22 | TVu-14195 | 1.35 | 37.71 | TVu-1780 | 1.45 | 12.00 | TVu-6968 | 1.31 | 28.35 | TVu-9820 | 0.99 | 27.76 |
| CC-110-1 | 0.66 | 30.80 | KVx-421-25 | 1.19 | 50.86 | TVu-14224 | 0.74 | 41.38 | TVu-1811 | 1.40 | 14.67 | TVu-7362_R | 0.94 | 12.92 | TVu-9848 | 1.39 | 32.00 |
| CC-36 | 1.37 | 42.67 | KVx-525 | 4.41 | 18.09 | TVu-14248 | 1.54 | 52.71 | TVu-1886 | 1.64 | 8.00 | TVu-7559 | 1.66 | 23.23 | TVu-9866 | 1.12 | 79.33 |
| CC-85-2 | 0.83 | 6.22 | KVx-61-1-1 | 1.54 | 45.87 | TVu-14253 | 1.99 | 50.67 | TVu-1916 | 1.37 | 212.57 | TVu-7625 | 0.84 | 12.16 | Tete-2 | 1.14 | 32.00 |
| CRSP_Niebe | 1.25 | 14.22 | Lori_Niebe | 1.00 | 126.00 | TVu-14272 | 1.88 | 134.00 | TVu-201 | 1.30 | 3.48 | TVu-7642 | 1.02 | 11.43 | UCR_11 | 0.83 | 35.20 |
| Cameroon_12-58 | 0.46 | 14.29 | Lyg_321-2 | 2.76 | 69.33 | TVu-14321 | 0.98 | 7.30 | TVu-21 | 0.63 | 13.79 | TVu-7647 | 1.23 | 23.77 | UCR_1340 | 1.48 | 18.53 |
| Cameroon_7-29 | 1.13 | 12.80 | Massava-11 | 1.74 | 118.00 | TVu-14336 | 1.37 | 33.94 | TVu-2168 | 2.94 | 24.20 | TVu-7684 | 1.15 | 7.00 | UCR_1342 | 2.41 | 56.53 |
| China_501 | 0.96 | 6.96 | Melakh | 1.82 | 47.00 | TVu-14345 | 5.74 | 149.33 | TVu-2280 | 0.89 | 19.00 | TVu-7719 | 1.07 | 6.00 | UCR_1432 | 1.21 | 70.96 |
| Cp_4877 | 1.32 | 15.06 | Montiero | 1.81 | 19.81 | TVu-14346 | 1.29 | 22.30 | TVu-2322 | 0.95 | 17.00 | TVu-7739 | 1.39 | 16.41 | UCR_162 | 1.28 | 12.00 |
| Cp_4906 | 1.42 | 22.74 | Mougne | 2.11 | 18.67 | TVu-14393 | 1.05 | 21.78 | TVu-2398 | 1.07 | 27.73 | TVu-7755 | 1.06 | 44.80 | UCR_193 | 1.68 | 17.33 |
| Cp_5556 | 1.86 | 31.38 | Mouride | 1.85 | 33.85 | TVu-14401 | 4.19 | 9.70 | TVu-2418 | 0.99 | 24.38 | TVu-7778 | 0.98 | 44.31 | UCR_24 | 1.24 | 41.00 |
| Cp_5647 | 1.51 | 32.94 | Moussa_Local | 2.03 | 19.83 | TVu-14533 | 0.93 | 15.11 | TVu-2449 | 1.31 | 49.23 | TVu-7798 | 1.46 | 28.80 | UCR_288 | 2.45 | 9.70 |
| Danila | 1.15 | 27.76 | Muinana-Lawe | 0.51 | 28.44 | TVu-14621 | 0.57 | 32.00 | TVu-2548 | 1.07 | 26.67 | TVu-7971 | 1.25 | 34.35 | UCR_3301 | 0.72 | 12.24 |
| Early_Scarlet | 1.07 | 26.67 | N'diambour | 1.20 | 34.13 | TVu-14632 | 1.19 | 37.87 | TVu-264 | 2.49 | 19.20 | TVu-7991 | 1.71 | 20.89 | UCR_3326 | 1.26 | 50.67 |
| Ecute | 1.20 | 9.78 | NamuesseD | 2.06 | 38.86 | TVu-14633 | 1.14 | 75.43 | TVu-2680 | 1.58 | 25.71 | TVu-8059 | 0.81 | 16.92 | UCR_5164 | 3.00 | 67.00 |
| FN-1-13-04 | 1.21 | 9.04 | Nhacoongo-1 | 1.41 | 14.40 | TVu-14691 | 1.47 | 53.94 | TVu-2723 | 2.74 | 96.00 | TVu-8072 | 1.57 | 13.63 | UCR_5272 | 1.33 | 21.76 |
| FN-1-14-04 | 0.76 | 61.82 | Nhacoongo-3 | 1.67 | 150.40 | TVu-14759 | 1.09 | 35.84 | TVu-2736 | 1.17 | 26.40 | TVu-8082 | 1.38 | 24.00 | UCR_5275 | 0.84 | 29.54 |
| FN-2-13-04 | 0.89 | 19.56 | Pakau3 | 1.21 | 63.48 | TVu-1477 | 1.08 | 40.00 | TVu-2845 | 1.44 | 21.33 | TVu-8262 | 1.18 | 20.57 | UCR_5279 | 1.40 | 24.00 |
| Faef-14-Inhaca-E | 4.62 | 17.89 | Prima | 1.46 | 36.00 | TVu-14862 | 1.44 | 51.39 | TVu-2933 | 3.22 | 20.00 | TVu-8389 | 1.39 | 9.38 | UCR_5280 | 1.01 | 90.67 |
| French_Bean | 0.79 | 34.53 | SASAQUE | 2.92 | 35.43 | TVu-14875 | 1.71 | 44.00 | TVu-2968 | 1.50 | 37.12 | TVu-84 | 1.23 | 20.29 | UCR_5353 | 0.74 | 82.91 |
| Gorda | 1.18 | 11.20 | Sanzi | 1.40 | 41.51 | TVu-14949 | 1.41 | 22.22 | TVu-2971 | 0.98 | 60.39 | TVu-8612 | 1.05 | 24.00 | UCR_5385 | 1.28 | 71.38 |
| IAR7/8-4-5-1 | 2.03 | 36.00 | Suvita-2 | 1.66 | 24.84 | TVu-14970 | 1.92 | 30.00 | TVu-3043 | 1.17 | 73.60 | TVu-8622 | 0.86 | 71.47 | UCR_707 | 1.61 | 80.00 |
| INIA-120_R | 1.10 | 11.10 | TVu-1000 | 1.94 | 12.80 | TVu-14971 | 1.71 | 10.67 | TVu-3076 | 1.88 | 34.56 | TVu-8631 | 1.79 | 7.47 | UCR_739 | 2.09 | 12.90 |
| INIA-31 | 1.34 | 34.67 | TVu-1004 | 1.33 | 32.00 | TVu-15114 | 0.99 | 76.00 | TVu-3156 | 2.38 | 18.76 | TVu-8641 | 1.37 | 19.64 | UCR_779 | 2.82 | 19.84 |
| INIA-34 | 1.48 | 15.36 | TVu-10100 | 1.75 | 41.71 | TVu-15143 | 1.25 | 25.78 | TVu-3282 | 2.33 | 107.43 | TVu-8656 | 1.21 | 13.54 | UCR_830 | 1.43 | 13.71 |
| INIA-40 | 1.03 | 27.33 | TVu-1016 | 0.76 | 28.44 | TVu-15315 | 0.86 | 26.46 | TVu-3360 | 1.30 | 1.43 | TVu-8671 | 0.71 | 25.45 | VAR-10B | 1.67 | 25.60 |
| INIA-41 | 0.91 | 7.00 | TVu-10179 | 0.72 | 42.86 | TVu-1536 | 1.71 | 56.00 | TVu-3552 | 1.41 | 48.40 | TVu-8673 | 1.38 | 36.00 | VAR-3A | 0.71 | 19.43 |
| INIA-5E | 1.09 | 24.00 | TVu-10281 | 2.64 | 22.77 | TVu-15391 | 1.48 | 54.40 | TVu-3565 | 1.59 | 75.00 | TVu-8702 | 0.88 | 32.52 | VAZZANO_Blk | 2.56 | 27.08 |
| INIA-72 | 1.19 | 50.46 | TVu-1036 | 1.47 | 24.00 | TVu-15400 | 2.35 | 104.30 | TVu-3652 | 2.00 | 34.00 | TVu-8713 | 1.97 | 51.81 | VAZZANO_Brw | 1.54 | 29.33 |
| IT00K-901-6 | 1.84 | 8.00 | TVu-10366 | 1.90 | 12.80 | TVu-15411 | 1.44 | 77.33 | TVu-3804 | 1.38 | 23.51 | TVu-8775 | 1.92 | 20.85 | Vg48 | 1.71 | 29.82 |
| IT82E-18 | 2.24 | 23.33 | TVu-1037 | 1.09 | 32.50 | TVu-15426 | 1.06 | 145.45 | TVu-3817 | 1.04 | 37.60 | TVu-8779 | 1.33 | 12.00 | Vg50 | 1.45 | 20.27 |
| IT83D-442 | 0.76 | 11.20 | TVu-10466 | 1.40 | 22.00 | TVu-15445 | 1.97 | 116.57 | TVu-3830 | 1.91 | 155.43 | TVu-8812 | 2.10 | 17.68 | Vg56 | 0.96 | 24.53 |
| IT84S-2049 | 0.83 | 16.94 | TVu-10513 | 2.24 | 42.88 | TVu-1556 | 2.33 | 116.87 | TVu-3842 | 1.15 | 105.29 | TVu-8834 | 1.72 | 39.11 | Vg58 | 1.56 | 18.82 |
| IT84S-2246 | 1.27 | 20.49 | TVu-1059 | 1.14 | 29.87 | TVu-15591 | 3.36 | 51.00 | TVu-393 | 1.38 | 41.85 | TVu-8877 | 1.44 | 15.30 | Vg62 | 1.42 | 18.82 |
| IT85F-867-5 | 1.07 | 28.57 | TVu-10892 | 1.11 | 33.88 | TVu-15610 | 1.86 | 66.46 | TVu-3947 | 3.21 | 26.18 | TVu-8883 | 1.47 | 13.84 | Vg72 | 1.13 | 25.33 |
| IT86D-364 | 0.58 | 14.15 | TVu-1124 | 1.79 | 54.00 | TVu-15636 | 1.13 | 45.41 | TVu-3965 | 4.51 | 9.41 | TVu-889 | 1.97 | 45.71 | Vg73 | 1.61 | 58.91 |
| IT89KD-288 | 1.15 | 15.56 | TVu-113 | 0.87 | 76.57 | TVu-15639 | 1.23 | 41.06 | TVu-4097 | 1.66 | 24.00 | TVu-8923 | 1.42 | 25.78 | Xingove | 1.23 | 11.50 |
| IT93K-452-1 | 1.14 | 16.59 | TVu-12565 | 2.63 | 29.87 | TVu-15653 | 2.57 | 10.43 | TVu-415 | 0.82 | 76.19 | TVu-8934 | 0.97 | 21.00 | Yacine | 1.71 | 12.80 |
| IT93K-503-1 | 0.80 | 46.93 | TVu-12710 | 1.64 | 9.50 | TVu-15661 | 0.77 | 24.83 | TVu-43 | 0.76 | 67.88 | TVu-9031 | 2.98 | 9.51 | | | |
| IT95K-1093-5 | 0.98 | 17.52 | TVu-12746 | 2.20 | 211.56 | TVu-15687 | 1.77 | 7.74 | TVu-4316 | 2.28 | 18.62 | TVu-9073 | 1.47 | 35.56 | | | |
| IT95K-1105-5 | 1.10 | 24.32 | TVu-1280 | 1.56 | 63.06 | TVu-1583 | 1.02 | 19.39 | TVu-4535 | 1.44 | 35.33 | TVu-9095 | 3.51 | 34.21 | | | |
| IT95K-1479 | 1.11 | 12.57 | TVu-12804 | 1.18 | 45.14 | TVu-15861 | 2.39 | 16.83 | TVu-4545 | 4.24 | 90.35 | TVu-9256 | 1.27 | 5.50 | | | |
| IT95K-1491 | 1.14 | 6.29 | TVu-12873 | 2.68 | 56.47 | TVu-15878 | 0.89 | 19.13 | TVu-4557 | 1.13 | 9.00 | TVu-9257 | 2.61 | 19.69 | | | |
| IT95M-190 | 0.99 | 7.36 | TVu-12897 | 2.00 | 103.43 | TVu-15973 | 1.18 | 12.00 | TVu-456 | 1.62 | 39.23 | TVu-9259 | 0.90 | 7.11 | | | |
| IT96D-610 | 1.41 | 5.12 | TVu-12923 | 2.68 | 108.44 | TVu-15995 | 0.84 | 6.40 | TVu-4622 | 2.06 | 68.21 | TVu-9265 | 1.31 | 27.29 | | | |
| IT97K-207-15 | 0.75 | 8.00 | TVu-12937 | 0.63 | 39.27 | TVu-1609 | 1.09 | 33.60 | TVu-4632 | 0.92 | 17.68 | TVu-9393 | 1.76 | 13.33 | | | |
| IT97K-207-21 | 1.02 | 7.25 | TVu-12968 | 8.55 | 30.40 | TVu-16220 | 0.82 | 46.40 | TVu-4669 | 1.44 | 46.12 | TVu-946 | 1.67 | 20.92 | | | |
| IT97K-461-4 | 1.05 | 8.43 | TVu-13017 | 1.75 | 29.71 | TVu-16237 | 0.97 | 40.89 | TVu-467 | 1.00 | 136.00 | TVu-9469 | 1.18 | 82.46 | | | |
| IT97K-499-35-1-1 | 0.71 | 31.04 | TVu-132 | 1.90 | 17.66 | TVu-16269 | 0.97 | 48.64 | TVu-4711 | 1.61 | 70.24 | TVu-9474 | 1.34 | 9.03 | | | |
| IT97K-499-39-1-1 | 1.42 | 16.30 | TVu-1330 | 1.25 | 12.31 | TVu-16278 | 1.04 | 11.64 | TVu-4984 | 1.16 | 23.32 | TVu-9486 | 1.40 | 14.19 | | | |
| IT97K-556-6 | 1.48 | 43.00 | TVu-13305 | 2.27 | 27.33 | TVu-16304 | 1.30 | 37.76 | TVu-53 | 4.17 | 58.67 | TVu-9492 | 1.58 | 47.33 | | | |
| IT97K-569-9 | 0.96 | 16.59 | TVu-13388 | 0.69 | 55.70 | TVu-16368 | 1.05 | 29.91 | TVu-5444 | 1.01 | 77.09 | TVu-9506 | 0.91 | 28.80 | | | |

**Table 1, Ethylene response ratios of 364 cowpea accessions belonging to the UCR Minicore.** Et/g ratios of *Vu*-In$^{-A}$ : $H_2O$ and *Vu*-In *:* H2O are shown. Cowpea trifoliate leaves were treated with 1 μM *Vu*-In or *Vu*-In$^{-A}$.

| | |
|---|---|
| AtSERK1 Fw | cacc ggtacc ATGGAGTCGAGTTATGTGGTGTT |
| AtSERK1 Rev | GTCA GCGGCCGC gc CCTTGGACCAGATAACTCAACGG |
| AtSERK2 Fw | cacc ggtacc ATGGGGAGAAAAAAGTTTGAAGCT |
| AtSERK2 Rev | GTCA GCGGCCGC gc TCTTGGACCAGACAACTCCATAG |
| AtSERK3 Fw | cacc ggtacc ATGGAACGAAGATTAATGATCC |
| AtSERK3 rev | GTCA GCGGCCGC gc TCTTGGACCCGAGGGGTATTC |
| AtSERK4 fw | cacc ggtacc ATGACAAGTTCAAAAATGGAACAAAG |
| AtSERK4 rev | GTCA GCGGCCGC gc TCTTGGACCCGAGGGGTAATCG |
| AtSOBIR fw | cacc ggtacc ATGGCTGTTCCCACGGGAAG |
| AtSOBIR rev | GTCA GCGGCCGC gc GTGCTTGATCTGGGACAACATG |
| Vigun09g096400 (VuSOBIR1) Fw | cacc ggtacc ATGGTTGCCAAAACTCACTTATTGCTTC |
| Vigun09g096400 (VuSOBIR1) Rev | GTCA GCGGCCGC gc GTGTTTGATCTGAGACAGCATGCAC |
| Glyma.10G227900 fw | cacc ggtacc ATGGATGCCTCTCCCTCAAGTATG |
| Glyma.10G227900 Rev | GTCA GCGGCCGC gc ATAGATGGAGTTCATCTTTTGGATCAC |
| Gm10 45840409 fw | AGTGGGAACTAAATACAGATTTTCC |
| Gm10 45843661 rev | TCCGAATAACACATTGGGCCT |
| Gm10 45854550 fw | TGCGCCTTTGCATTATTCCAC |
| Gm10 45862505 rev | AGCAAAAGAGCCCAGAGACC |
| INR-Vu Fw | cacc ggtacc ATGGATGCCTTCAAACTCTTTCTAC |
| INR-Vu Rev | GTCA GCGGCCGC gc ATGGATGTAAGCACGGTTGCATC |
| Vigun219700 Fw | cacc ggtacc ATGTGCCATTATAAATACTTCCCATGG |
| Vigun219700 Rev | GTCA GCGGCCGC gc CATTCCAGTAATGGAGTTTGTCTTTTGG |
| Phvul.007G077500 Fw | cacc ggtacc ATGAATGCCTCTCCCTTCAATATTGAAC |
| Phvul.007G077500 Rev | GTCA GCGGCCGC gc ATGCATGTAAGCATGGTTGCATC |
| Vradi08g18330 Fw | cacc ggtacc ATGGATGCCTTCAGCATTGAACTC |
| Vradi08g18330 Rev | GTCA GCGGCCGC gc ATGGATGTAGGCACGGTTGCATC |

**Table 2, Primers used in the study for gene amplification.** 5’ to 3’ sequences shown

**4. References 27-43.**

27. W. Muchero et al., A consensus genetic map of cowpea [Vigna unguiculata (L) Walp.] and synteny based on EST-derived SNPs. Proc. Natl. Acad. Sci. U. S. A. 106, 18159–18164 (2009).

28. Y. Wu, P. R. Bhat, T. J. Close, S. Lonardi, Efficient and accurate construction of genetic linkage maps from the minimum spanning tree of a graph. PLoS Genet. 4, e1000212 (2008).
